# Context-dependent effects of ivermectin residues on dung insects: Interactions with environmental stressors, size, and sex in a sepsid fly (*Sepsis neocynipsea*)

**DOI:** 10.1101/2024.11.18.623968

**Authors:** Jill Walker, Benjamin J. Mathews, Patrick T. Rohner

**Author notes:** Shared first co-authorship (Jill Walker, Benjamin J. Mathews). Corresponding Author: Department of Ecology, Behavior, and Evolution, University of California San Diego, La Jolla.

## Abstract

Insects associated with livestock dung frequently encounter veterinary medication residues. These residues often have negative effects on insect survival, reproduction, and ecosystem functioning and may contribute to the rapid decline in temperate insect populations. Ivermectin is an antiparasitic drug widely used to treat parasites in livestock. While it has long been recognized that ivermectin effects insect survival, the potential interactive effects between ivermectin exposure and other ecologically relevant abiotic stressors remain poorly understood. Here, we study these effects in the black scavenger fly *Sepsis neocynipsea*, which depends on cow dung for reproduction. Using a fully factorial design, we test whether the effects of ivermectin exposure on adult survival interact with heat and desiccation stress, and whether these effects depend on size and sex. Ivermectin exposure had strong negative impacts on adult survival, but its effects were stronger in females and large individuals. While heat stress also had a strong effect on adult survival, the combined effects of heat and ivermectin exposure were less severe than the expected additive effects of both stressors applied independently, suggesting some cross-resistance. We did not find an interaction between ivermectin and desiccation stress. Taken together, our findings highlight how the complex interactions between insecticides and abiotic stressors could drive changes in coprophagous insect populations and their ecological functions across different ecosystems and climates.

## Introduction

The environmental impacts of agricultural chemicals are a major concern in conservation and environmental management (Zhou et al. 2025). Antiparasitic drugs are a large component of the animal health market accounting for €7 billion in annual sales as of 2018 (Selzer and Epe 2021). These drugs treat an increasingly broad-spectrum of endo- and ectoparasites and greatly benefit animal and human health (Crump 2017). Despite these benefits, there have been concerns over the ecological impact of antiparasitic drugs when they enter ecosystems. Even low doses of broad-spectrum antiparasitics can negatively affect off-target organisms in the environment (Gandara et al. 2024). This can be especially problematic for coprophagous (i.e., dung-eating) insects that are often in contact with chemical residues found in the feces of treated livestock.

The impact of antiparasitic residues has been heavily studied in coprophagous insects. This ecological guild includes members of several families of dung beetles and dung flies, and plays important roles in regulating the decomposition of fecal matter, especially in agricultural contexts (Kavanaugh and Manning 2020; Losey and Vaughan 2006; Skidmore 1991). Through consuming, burying, aerating, and fragmenting dung, coprophagous insects directly and indirectly facilitate fecal decomposition, often driving the local microbial, fungal, and invertebrate diversity (Stevenson and Dindal 1987). However, the reliance of coprophagous insects on dung as a food source frequently exposes them to chemicals secreted in vertebrate dung. One such chemical is ivermectin — an broad-spectrum antiparasitic drug often used to treat nematode, mite, and lice infections in livestock, pets, and humans (Crump 2017). As much as 80-90% of an ivermectin dose can be excreted through feces and, due to its chemical stability, residues can remain in the environment for weeks (Alvinerie et al. 1999; Herd et al. 1996). Because ivermectin is a broad-spectrum antiparasitic that acts on a wide range of arthropods and nematodes (El-Saber Batiha et al. 2020; McKellar 1997; Puniamoorthy et al. 2014), environmental residues can negatively impact the community of coprophagous invertebrates that rely on livestock feces for food and reproduction, thereby greatly impairing ecosystem functioning (Correa et al. 2022; Jochmann and Blanckenhorn 2016; Kavanaugh and Manning 2020; Verdú et al. 2018).

While ivermectin has been shown to be directly fatal to coprophagous flies and dung beetles through its effects on molting, growth, and reproduction (McKellar 1997; Pérez-Cogollo et al. 2015; Puniamoorthy et al. 2014; Rodríguez-Vivas et al. 2020; van Koppenhagen et al. 2020), exposure to ivermectin (and other avermectins) is not universally fatal to coprophagous insects (Conforti et al. 2018; Jochmann and Blanckenhorn 2016; Schmidt 1983). Recent studies also suggest that drug exposure might interact with heat stress (Bueno et al. 2023; González-Tokman et al. 2022; Sirois-Delisle and Kerr 2022) but the degree to which these context-dependent effects of ivermectin are common is unclear. Similarly, whether the negative effects relate to an insect’s intrinsic features, such as its size or sex (patterns that have been found in other species and other drugs, e.g.,: Rathman et al. 1992; Zhang et al. 2019), remains poorly understood. We here test these effects in black scavenger flies.

Black scavenger flies (Sepsidae) are a functionally important group of insects highly abundant in temperate and alpine grasslands. Despite their small size, ranging from 2 to 6 mm in body length, they play key roles as detritivores, pollinators, and a food source for other invertebrates (Pont and Meier 2002; Rohner et al. 2014; Rohner et al. 2019). Most sepsid species rely on vertebrate excrement for reproduction, with adult females of many species laying their eggs on the surface of fresh cow dung. The larvae hatch and feed on the dung until they pupate either inside or near the cow pat. Cow dung is essential not only as a site for oviposition but also as a crucial food source for egg production in females (Pont & Meier 2002). Given their ecological role and dependence on vertebrate dung across their entire lifecycle, sepsids are an ideal system for studying the ecosystem effects of veterinary pharmaceuticals and their context dependency.

Previous studies in sepsid flies have shown that sepsid larvae are very sensitive to ivermectin. Larval sepsids exposed to drug residues in cow dung show high mortality, even at relatively low and ecologically relevant concentrations (Blanckenhorn et al. 2013; Puniamoorthy et al. 2014). Adult flies, which frequently visit dung pats for oviposition, mating, and feeding, are also significantly affected by ivermectin exposure, which can reduce their survival, fecundity, and fertility (Conforti et al. 2018). While large species differences in sensitivity to ivermectin have been documented, little is known about how ivermectin exposure affects individuals within a species under varying environmental conditions. In the yellow dung fly, recent studies suggest that exposure to high temperatures might exacerbate the negative effects of ivermectin residues (González-Tokman et al. 2022). However, whether such interactive effects are widespread is still unclear (Halsch et al. 2023). In addition to external environmental stressors, the effects of ivermectin exposure might also depend on an individual’s size or sex. A large body of research demonstrates that female insects are often more sensitive to nutritional conditions, possibly due to the costly development of eggs and ovaries (Rohner et al. 2018; Stillwell et al. 2010; Teder and Tammaru 2005). Similar effects could be expected for the exposure of chemical residues in the environment. Given female’s role in population growth rates, sex-specific effects could further exacerbate (or reduce) the ecological effects of ivermectin exposure. We here investigate how ivermectin exposure interacts with other external and internal factors in *Sepsis neocynipsea* — a species common in North American grasslands. Using a fully factorial design, we test for interactive effects between ivermectin exposure, heat and desiccation stress, as well as size- and sex-specific effects. We expected to observe strong synergistic effects between ecological and pharmaceutical stressors. In congruence with earlier studies, we find strong effects of ivermectin exposure on adult lifespan, but we also find that these effects are context dependent. This highlights previously overlooked dynamics that are likely to have important effects on ecosystem functioning.

## Methods

To assess the effects of veterinary antibiotic residues on coprophagous insects, we studied the effects of ivermectin exposure in *Sepsis neocynipsea* Melander and Spuler, 1917. This species has a broad Holarctic distribution and is one of the most abundant sepsid flies in North American agricultural landscapes (Pont and Meier 2002). We collected 30 females in Bloomington, Indiana, USA and used their offspring to establish large outbred laboratory colonies kept under standard conditions (see e.g., (Rohner et al. 2016)).

### Experimental design

To assess the context-dependence of ivermectin exposure on adult survival, we conducted replicated laboratory trials with fully factorial treatment combinations incorporating ivermectin treatments, heat stress, and desiccation stress. Each replicate trial consisted of a 1.9L plastic container equipped with a small 30 ml ramekin (deli cup) containing ca. 1.5 grams of sugar. 20 adult flies that were at least two weeks old were randomly assigned to each container.

To expose adult flies to an ivermectin treatment and a control, we added a 59ml cup filled with previously frozen cow dung to each replicate container. To assess the overall effect of ivermectin on adult survival, we either mixed 500µl acetone into the dung (as a control treatment), or 500µl of acetone containing 3 µg of ivermectin (leading to a total ivermectin concentration of 150µg of ivermectin kg^-1^ of dung (wet weight)). This concentration is on the low end of many field estimates of fecal ivermectin concentration (Fernandez et al. 2009; Lumaret et al. 2007), although predicted environmental concentrations vary greatly (Liebig et al. 2010). For both treatments, we allowed the acetone (which is used as solvent) to evaporate from the dung for five hours before collecting flies to place into their respective containers.

This ivermectin treatment was crossed with a desiccation stress treatment. Half of all experimental replicate containers included a 30 ml lidded plastic cup filled with water. We threaded a cotton string through the lid of the cup to draw water from the cup to the surface to make it available to the flies. The other half of all containers did not contain a water source. In those containers that contained a water cup, flies could access water from the water cup and the cup filled with dung. In the desiccation treatment, dung represented the only source of moisture.

In addition to the ivermectin and desiccation treatments, we also exposed half of all replicates to temperature stress. Half of all replicates were placed in an incubator calibrated to constant 23 degrees Celsius, while the other half were placed in an incubator calibrated to 33 degrees Celsius.

In total, each treatment combination had three experimental replicates, leading to a total of 482 flies. The experiment was run in two different temporal blocks. The first block contained one replicate per treatment combination while the second block contained two replicate containers.

### Estimating adult survival

We collected dead flies every 24hrs to record mortality and sexed each individual based on primary and secondary sexual morphology (Baur et al. 2019). The experiment was terminated when there were signs of late-stage pupae in the control treatments. To generate an estimate of overall body size, we removed the left and right wing of all individuals, embedded them in glycerol on a glass slide, and photographed them using a Pixelink camera (M20C-CYL) mounted on a Leica M205 stereoscope. We then used ImageJ to measure the length of the second longitudinal wing vein as an estimate of adult body size (See fig S1). Due to wing wear, size could not be estimated for all individuals. When measurements for the left and right side were available, we used the mean for further analysis.

### Statistical analyses

The effects of ivermectin exposure, sex, body size, temperature, desiccation stress, as well as all interactions on adult survival were analyzed using a Cox mixed-effects model using maximum likelihood as implemented in the R-package ‘coxme’ (Therneau 2022). The experimental replicate was added as a random effect. Individuals that were alive at the end of the experiment were entered as censored data. Wing length (see fig S1) was measured as an estimate of body size. Wing length was mean centered prior to the analysis. We started by fitting a full model with all possible interactions and subsequently removed all non-significant interaction terms.

## Results

To assess the effect of ivermectin exposure on adult survival, we exposed a total of 482 individual sepsid flies to six different treatment combinations. Across all treatments, 48.8 percent (235/482) of all individuals died in the course of the experiment.

To investigate the overall effect of ivermectin on adult survival, we first considered a model with ivermectin exposure as the sole dependent variable. This analysis revealed that exposure to ivermectin increased mortality risk more than 13-fold compared to the control treatment (Cox mixed-effects model with container as random effect: HR = 13.20, z = 4.05, P <.001). However, more complex models that included all treatments, sex, as well as size information revealed that the impact of ivermectin exposure on survival is context dependent.

The effect of ivermectin on adult survival was sex specific. Females were about three times less likely to survive compared to males (sex-by-ivermectin treatment interaction: HR = 0.33, z = - 3.00, P = 0.003, Table 1). The detrimental effects of ivermectin on survival were also reduced when exposed to high temperatures, a finding that goes against patterns found in other species (González-Tokman et al. 2022).

**Table 1:** Cox mixed-effects model fit by maximum likelihood (n = 466; 16 individuals with missing wing length excluded). (SE = standard error, HR = hazard ratio)

|  | **coefficient** | **SE** | **HR** | **Z** | **P** |
| --- | --- | --- | --- | --- | --- |
| Wing length (mean-centered) | 1.67 | 1.00 | 5.33 | 1.67 | 0.095 |
| Heat stress [23°C → 32°C] | 1.21 | 0.67 | 3.35 | 1.80 | 0.072 |
| Desiccation stress [control → desiccation] | 0.01 | 0.46 | 1.01 | 0.03 | 0.980 |
| Ivermectin treatment [control → antibiotic exposure] | 4.32 | 0.62 | 74.85 | 7.00 | **<.001** |
| Sex [female → male] | 0.12 | 0.32 | 1.13 | 0.38 | 0.700 |
| Wing length × Heat stress | -2.94 | 0.76 | 0.05 | -3.87 | **<.001** |
| Wing length × Desiccation stress | 1.61 | 0.72 | 5.03 | 2.23 | **0.025** |
| Wing length × Ivermectin treatment | -2.01 | 0.80 | 0.13 | -2.53 | **0.012** |
| Heat stress × Ivermectin treatment | -1.58 | 0.70 | 0.21 | -2.27 | **0.023** |
| Ivermectin treatment × Sex | -1.09 | 0.36 | 0.33 | -3.00 | **0.003** |
| Heat stress × Desiccation stress | 2.47 | 0.61 | 11.87 | 4.08 | **<.001** |

Effects of the various chemical and environmental stressors also depended on body size. Large individuals were more resistant towards high temperatures and ivermectin exposure compared to smaller individuals (size-by-ivermectin interaction: HR = 0.13, z = −2.53, P = 0.012; size-by-temperature interaction: HR = 0.05, z = −3.87, P = 0.001, Table 1). However, larger individuals were more strongly affected by desiccation stress (size-by-desiccation interaction: HR = 5.03, z = 2.23, P = 0.025, Table 1). Larger individuals did therefore not have a survival advantage in all contexts. Desiccation and heat stress had very strong negative effects on survival when combined simultaneously (desiccation-by-temperature stress interaction: HR = 11.87, z = 4.08, P <.001, Table 1).

## Discussion

Dung insects are frequently exposed to chemical stress (Gandara et al. 2024), but the degree to which this interacts with other ecological stressors remains unclear. We investigated how ivermectin exposure, heat, and desiccation stress impacted the survival in adults of the sepsid fly *Sepsis neocynipsea*. Our results indicate that ivermectin exposure significantly increases mortality, though this effect varied by context. Mortality was notably higher among females and smaller individuals, while the lethal effects of ivermectin depended on temperature. Larger flies showed greater resilience to heat stress but were more vulnerable to desiccation. Overall, complex interactions among sex, temperature, desiccation, and ivermectin exposure can—depending on the specific conditions—modulate the mortality rate caused by ivermectin exposure. These findings have important implications for the ecology of black scavenger flies in agricultural systems, particularly as these ecosystems face increasingly strong environmental change.

### Interactions between different ecological stressors in complex environments

In natural environments, insects are exposed to varied, interacting stressors that are often not captured in experiments focused on exposure to singular stressors (Bueno et al. 2023; Rodrigues and Beldade 2020; Rohner and Moczek 2023). Interactions between chemical residues and other environmental stresses could contribute to the rapid decline of insect populations observed in the field (e.g., Wagner et al. 2021). Previous research has shown that heat stress and ivermectin exposure can have negatively synergistic effects on offspring survival in yellow dung flies (González-Tokman et al. 2022), a distantly related group of flies that is also dependent on cow dung for reproduction. Similarly, we expected heat and desiccation stress to greatly increase mortality in sepsid flies. While our results indeed revealed strong interactive effects between heat and desiccation (see Fig. 1), we did not find the expected interactions between ivermectin exposure and either of these major abiotic stressors. Specifically, desiccation stress did not interact with ivermectin, and the relationship between ivermectin and heat stress proved more complex than anticipated. Although ivermectin reduced survival when combined with temperature stress compared to temperature stress alone, this effect was much weaker than the impact of ivermectin at a control temperature, leading to a significant interaction term. This unexpected finding may indicate a form of cross-resistance, where exposure to one stressor confers increased resistance to another. Temperature has previously been shown to improve insecticide resistance in various insect pests (Bueno et al. 2023). Heat hardening—a physiological adaptation that enhances tolerance to high temperatures (Gu et al. 2019; Moghadam et al. 2019; Sørensen et al. 2019)—might represent a potential mechanism through which sepsids become more resistant against ivermectin through behavioral or physiological mechanisms. If so, the ancient adaptive response to heat could serve as an exaptation for handling exposure to ivermectin, which has only recently become widespread in agricultural environments. Overall, these findings suggest that examining the interactive effects of ivermectin and other environmental stressors is essential to understanding antibiotic impacts in dynamic and rapidly changing ecosystems.

**Figure 1:**
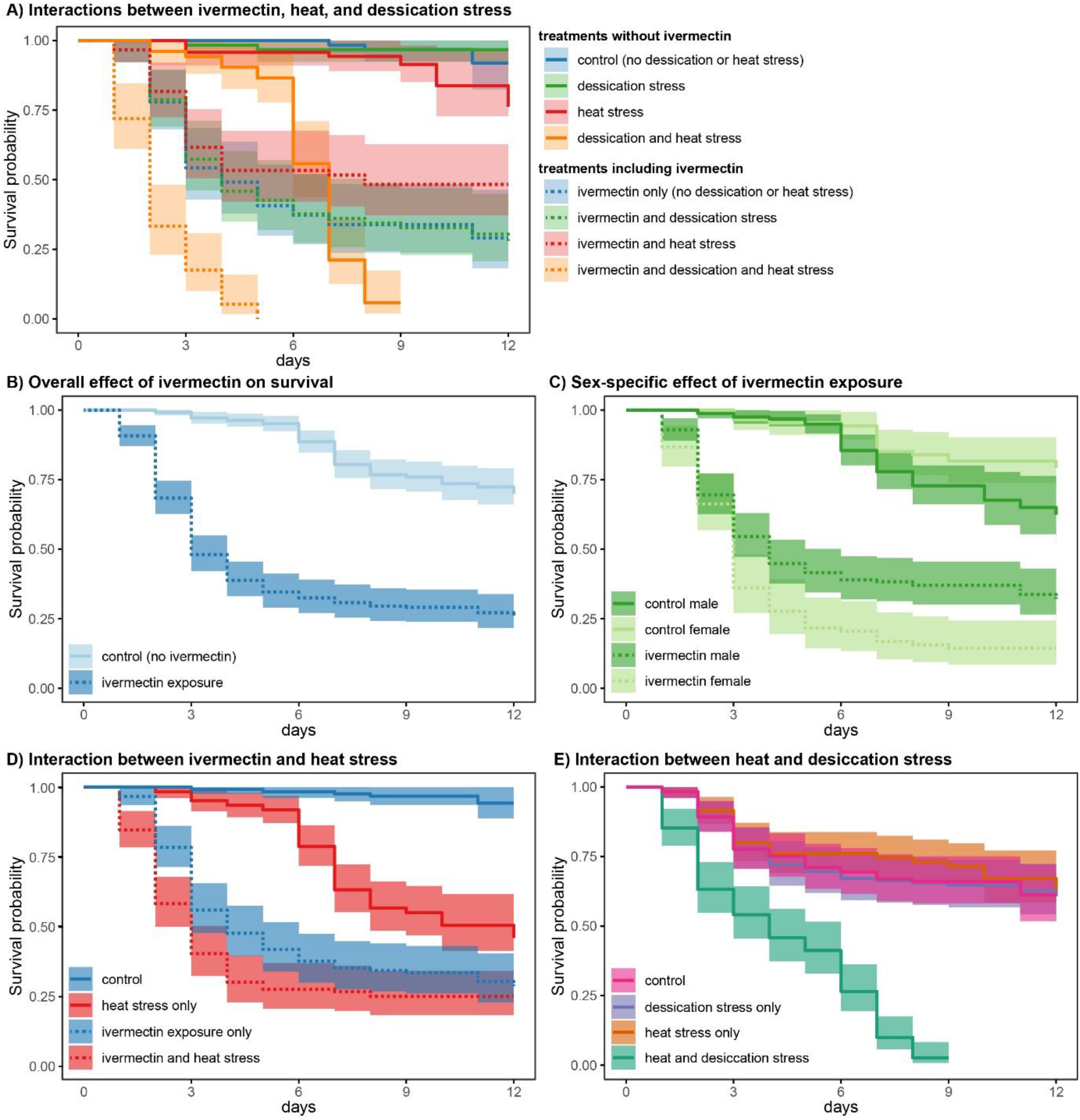
Effect of ivermectin in combination with other ecological stressors on adult survival of *Sepsis neocynipsea*. Plots show survival probability with time and associated 95% confidence limits. Panel A shows the combined effects of desiccation and heat stress in combination with ivermectin exposure (hatched line). Panels B-E highlight some of the key effects and interactions found in the overall model.

### Sex-specific effects of ivermectin exposure on adult survival

Our findings indicate that females are about three times more likely to die from ivermectin exposure than males. These sex-specific effects of ivermectin are a possible reflection of the disproportionate nutritional needs of females. In many insect species, including blowflies (Vogt et al. 1985), females must feed extensively to support egg production, which requires more protein-rich nutrients than the nectar that males typically consume. Because sepsid flies do not seem able to differentiate between contaminated and ivermectin-free cow dung (Conforti et al. 2018), this would lead to an increased exposure to ivermectin in females. This could possibly cause the sex difference in mortality when exposed to the same contaminated environment.

Alternatively, females may be more sensitive to chemical stresses due to their physiology. Previous studies indicate that females are often more vulnerable to nutrient stress than males (Teder and Tammaru 2005), and similar patterns could extend to chemical stress, although higher female mortality is not consistent across insect groups (e.g., Andreazza et al. 2020). Regardless of the underlying mechanism, increased female mortality is concerning, as the number of females in a population determines its potential growth rate. Ivermectin-driven reductions in female numbers could therefore significantly impact population dynamics and genetic diversity (Sutton et al. 2014). Further research is needed to determine whether these sex-specific effects are widespread.

### Interactions between ivermectin exposure and body size

We found that larger individuals had a reduced mortality when exposed to ivermectin and greater resistance to high temperatures. Larger individual’s resistance to high temperatures aligns with the findings of previous studies showing that larger individual insects have greater heat tolerance (e.g., Baudier et al. 2015). Large size thus seems to provide fitness benefits in terms of increased survival. However, large individuals were also more strongly affected by desiccation stress. The latter conflicts with the findings of several studies in other systems suggesting that large individuals are more resistant to desiccation stress (Bujan et al. 2016; Chown and Nicolson 2004; Hadley 1994). One possibility is that behavioral responses and microhabitat choice are the main mechanisms that regulate desiccation stress in sepsids (as has been shown in other systems, e.g., Hood and Tschinkel 1990). Because our experimental setup limited behavioral responses, it is unclear whether the physiological responses that are detected under laboratory conditions are relevant in the field. Future research will be necessary to investigate the interactions between desiccation, plastic life history responses, and behavior under more natural conditions.

## Conclusions

Antibiotic residues have long been recognized to negatively affect insect physiology, reproduction, and ecosystem functions of dung insects. Our data suggest that previous research have overlooked important interactions with sex, body size, and other ecological stressors that are likely to exacerbate the ecological impact of ivermectin (but see: González-Tokman et al. 2022). Moreover, most studies on the effects of ivermectin exposure have focused on temperate insect species. Considering the high insect diversity in equatorial regions and the rapid warming in Arctic environments, future research should prioritize understanding the interactions between abiotic stressors and ivermectin exposure in dung fly species from both equatorial and Arctic ecosystems.

## Data Availability Statement

The data that support the findings of this study are deposited in Dryad at datadryad.org: http://datadryad.org/stash/share/OWntylDqTP0cvgck4mx9y21zpmfyExN7kc2rw56p5zk

## Supplementary Material

**Figure S1:**
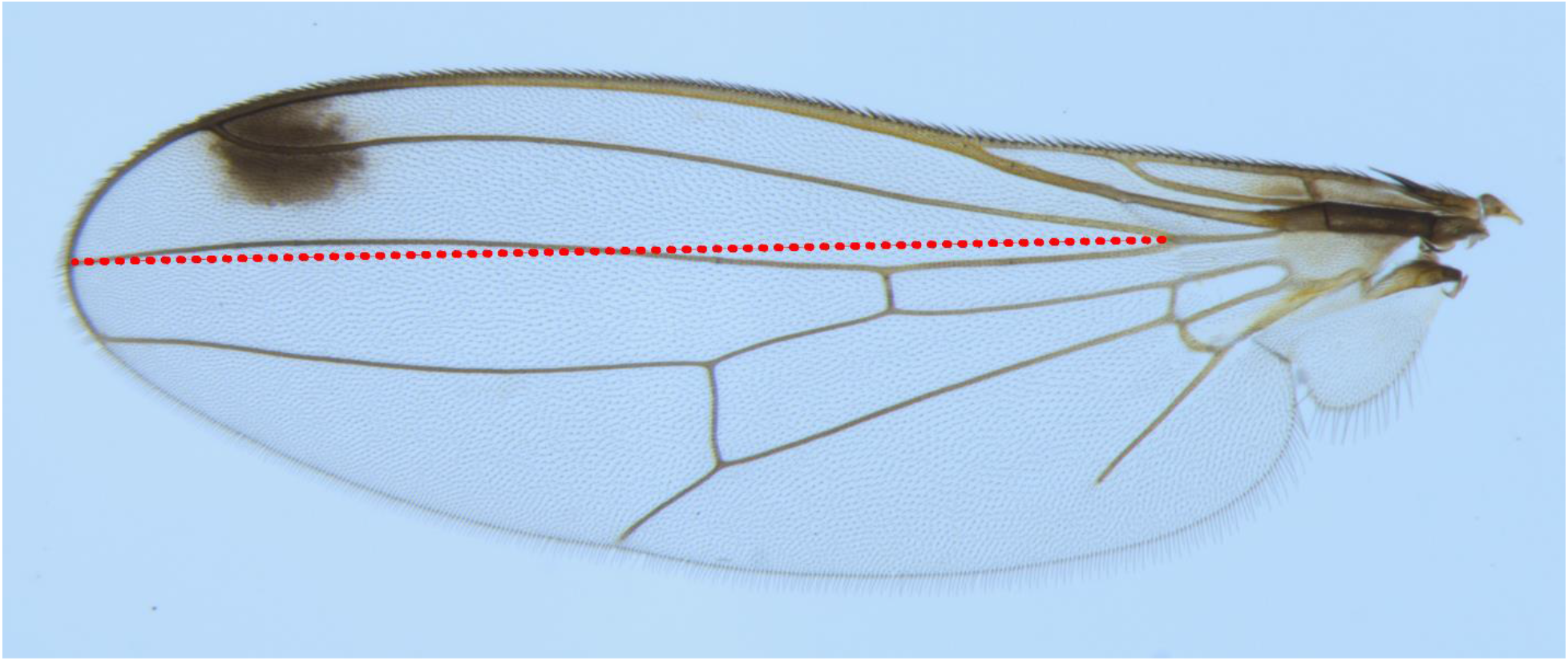
The points on the wing of *Sepsis neocynipsea* used to indicate body size. The wing of individual 266, a female sepsid fly exposed to ivermectin, desiccation stress, and the low temperature condition (23 degrees Celsius) is pictured to exemplify how wing length was measured.

